# BAG6 and RNF126 are broadly involved in protein quality control of non-native missense protein variants

**DOI:** 10.64898/2026.02.04.703735

**Authors:** Line Pedersen, Jonas Dideriksen, Kristoffer E. Johansson, Celeste M. Hackney, Isa K. Henrichs, Erna S. Sigmarsdóttir, Martin Grønbæk-Thygesen, Vasileios Voutsinos, Kresten Lindorff-Larsen, Rasmus Hartmann-Petersen

**Affiliations:** Department of Biology, University of Copenhagen, Ole Maaløes Vej 5, 2200N Copenhagen, Denmark

**Keywords:** ubiquitin, proteasome, UPS, chaperone, proteostasis, misfolding, Parkinsons disease, phenylketonuria, PKU, folliculin, Birt-Hogg-Dubé, PRKN, HSP70, HSP90

## Abstract

Protein quality control (PQC) degradation is a vigorous and selective system that limits the accumulation of harmful, non-native and aggregation-prone proteins. Specific PQC pathways have been described for misfolded ER proteins, aberrant translational products and mislocalized proteins. However, many components involved in PQC of structurally unstable cytosolic proteins are unknown despite such proteins being a common consequence of pathogenic coding variants. Here, we present results from a genome-wide CRISPR knockout screen to identify components involved in PQC of the Parkin R42P variant. We find that HSP90 and STIP1 stabilize Parkin, while the BAG6 chaperone and E3 ubiquitin ligase RNF126 are critical for PQC degradation. Variant abundance by massively parallel sequencing (VAMP-seq) reveal >1000 Parkin variants, including several variants linked to autosomal recessive juvenile Parkinsonism, are BAG6 targets. Pathogenic missense variants in phenylalanine hydroxylase (PAH) and the tumor suppressor FLCN are also stabilized in BAG6 and RNF126 knockout cells. We propose that BAG6 and RNF126 are broadly involved in PQC and may represent targets for the development of therapeutics for protein misfolding diseases.

## Introduction

The function of most proteins depends on their ability to fold into a native structure. When folding fails, or if the native state is unstable, the resulting non-native proteins are prone to form toxic aggregates and engage in non-specific interactions with other cell components and therefore represent a significant threat to the cell. Consequently, in all cells, protein folding is carefully monitored by the protein quality control (PQC) system that relies on both chaperones (1-4) and specific components of the ubiquitin-proteasome system (UPS) (5). Molecular chaperones catalyze the folding of nascent proteins and refolding of misfolded proteins while E3 ubiquitin-protein ligases, often operating in conjunction with chaperones, target non-native proteins for proteasomal degradation (5). Autophagy also plays a role in PQC degradation but typically targets larger protein inclusions (6, 7).

Molecular genetics studies have linked many chaperones and co-chaperones to PQC degradation, including the major chaperones involved in protein folding, HSP70 and HSP90 (2, 8, 9). However, only a few E3 ubiquitin ligases have been directly connected with PQC degradation (5). The PQC E3s are likely to display broad substrate specificity, ensuring that any non-native protein can be recognized while leaving native proteins untouched. The distinguishing features of non-native proteins that are recognized by PQC E3s, the so-called PQC degradation signals (degrons), appear to be stretches of hydrophobic residues that are buried in the native conformation but become exposed upon unfolding or misfolding (10-14). Since such hydrophobic regions overlap with chaperone binding sites (15, 16), a close connection between chaperones and the PQC E3s is expected. This is exemplified by the E3 ubiquitin ligase CHIP (STUB1), which binds directly to chaperones allowing for ubiquitylation and subsequent degradation of the chaperone substrate (17, 18). Another example is the chaperone-E3 complex BAG6:GET4:UBL4A:RNF126 connected to the degradation of readthrough proteins (19), tail-anchored proteins (20, 21), extracted membrane proteins (22) and mislocalized proteins (23, 24).

Most proteins are only marginally stable in their natural environment. It is well established that a common pathogenic mechanism in human genetic disorders is missense variants leading to reduced structural stability of the encoded protein (25, 26). Indeed, recent computational modelling of all possible missense variants in all human proteins indicates that ∼40% of all missense variants that lead to disease likely do so by reducing protein stability (27). Reduced structural stability often results in PQC degradation, and thus, many disease-linked protein variants display low steady-state protein levels (25, 28). Importantly, pathogenic gene variants that reduce protein stability may not necessarily completely abolish protein function but will often simply reduce activity resulting in hypomorphic variants (26). Well-characterized examples of this include the cystic fibrosis CFTR F508Δ variant (29) and the Parkin R42P variant linked to autosomal recessive juvenile Parkinson’s disease (30, 31). Hence, a typical mechanism which causes pathogenic dysfunction involves the degradation of structurally destabilized yet functional protein variants, leading to insufficient cellular levels. In such cases, the loss-of-function phenotypes could, in principle, be rectified by boosting synthesis or blocking PQC degradation (25, 32). Identifying the components and characterizing the pathways involved in PQC are therefore a priority not only for furthering our understanding of proteostasis but also because the PQC system represents a potential drug target for a wide range of genetic disorders (25).

Here, we applied genome-wide CRISPR knockout screens combined with fluorescence-activated cell sorting to identify components involved in the turnover of the Parkin R42P variant. We find that BAG6 and RNF126 are critical for R42P turnover while HSP90 protects Parkin from degradation. Using a site-saturated library of Parkin variants, we show that BAG6 and RNF126 regulate the abundance of most non-native Parkin variants. Since we find that BAG6 and RNF126 also regulate the degradation of unrelated disease-linked protein variants, BAG6 and RNF126 appear to be broadly involved in PQC degradation of structurally destabilized missense proteins and may represent targets for the development of therapeutics to misfolding diseases.

## Results

### Parkin R42P is a rapidly degraded PQC target

To identify components involved in PQC degradation of structurally unstable missense protein variants, we selected the Parkin R42P variant, linked to autosomal recessive juvenile Parkinson’s disease (OMIM: 600116), as a model protein (33). Previous studies have shown that the R42P variant is unfolded, rapidly degraded, and present at a reduced steady-state level (31, 34-36). Wild-type Parkin and the R42P variant were expressed as GFP-fusion proteins from a genomic landing pad in human HEK293T cells (31, 37). To correct for cell-to-cell variations in expression, mCherry is produced from the same mRNA via an internal ribosomal entry site (IRES) (**Fig. 1A**). In agreement with our previous observations using this expression system (31), western blotting revealed that the R42P steady-state level was reduced (**Fig. 1B**). This reduction was also evident by flow cytometry (**Fig. 1C,D**). Since the R42P level increased in response to inhibition of the proteasome and the ubiquitin E1, using bortezomib (BZ) and TAK243 (MLN7243), respectively (**Fig. 1C,D**), we conclude that the reduced steady-state level is caused by ubiquitin-dependent proteasomal degradation. Finally, we noted that upon blocking the molecular chaperone HSP90 using geldanamycin, Parkin abundance was reduced (**Supplementary Fig. 1**), indicating that HSP90 contributes to the folding and stability of the protein. We conclude that Parkin R42P is targeted for ubiquitin- and proteasome-dependent PQC degradation.

**Fig. 1.**
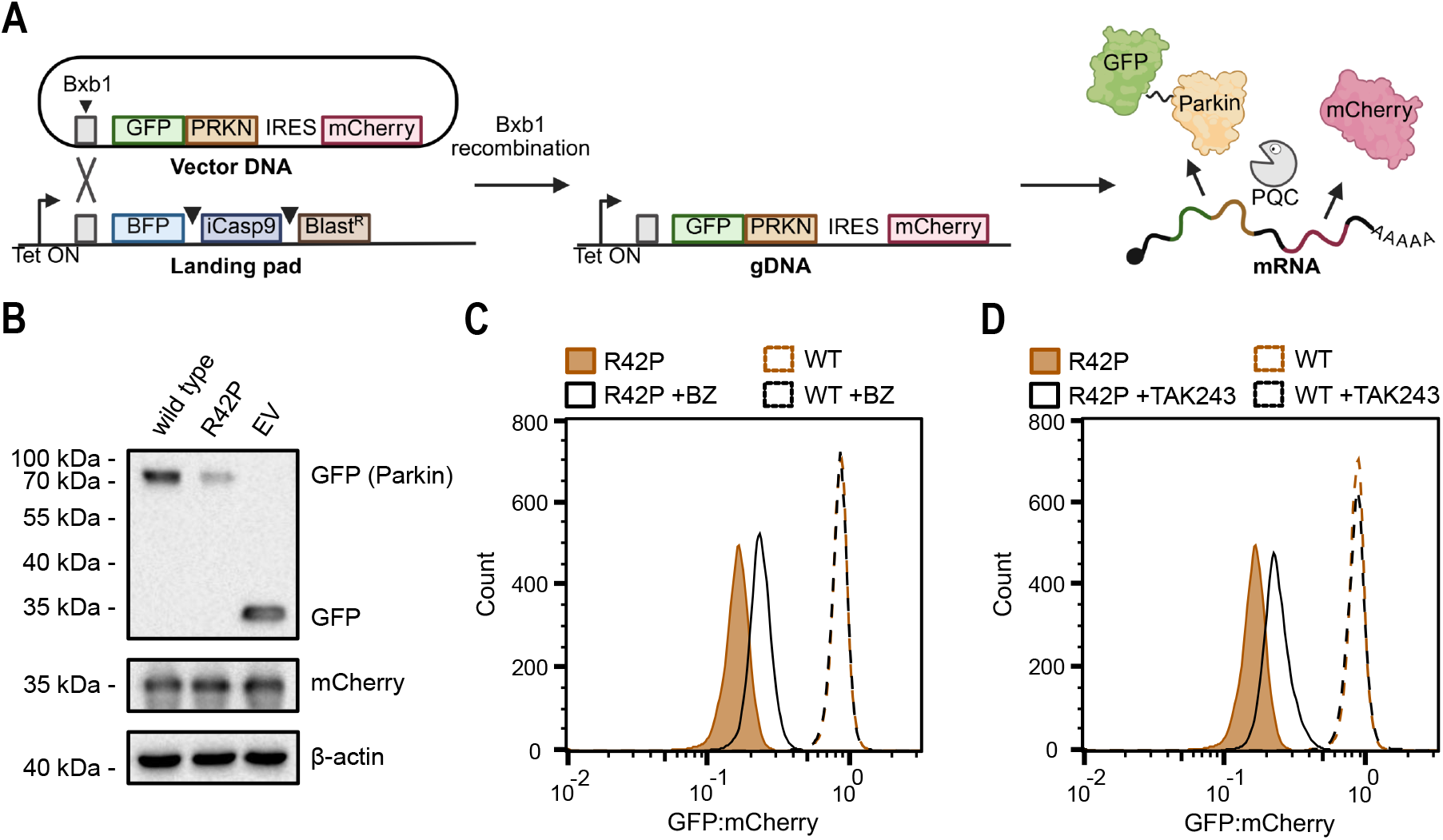
Parkin R42P is a target of the ubiquitin-proteasome system. (A) Schematic illustration of the expression system. For single copy expression, a plasmid (vector DNA) encoding GFP-Parkin followed by an internal ribosomal entry site (IRES) and mCherry, was introduced into the landing pad in the HEK293T genome (gDNA) by Bxb1 site-specific recombination. This displaces the cassette of BFP-iCasp9-BlastR separated by 2A stop/start sites (arrowheads). The expressed GFP-tagged Parkin protein variant is subject to PQC degradation (Pacman) resulting in reduced GFP fluorescence, while the mCherry level is constant. Created in BioRender. Hartmann-Petersen, R. (2025) https://BioRender.com/m5by8sy. (B) Comparison of protein levels of wild type Parkin, the R42P variant and empty vector (EV) by SDS-PAGE and western blotting of whole cell lysates. mCherry and β-actin were included as controls. (C) Representative flow cytometry profiles for landing pad cells expressing Parkin wild-type (n = 9.8×10^3^) and R42P (n = 1×10^4^) treated with 10 µM bortezomib (BZ) for 16 hours. Untreated wild-type (n = 9.7×10^3^) and R42P (n = 1×10^4^) cells are included for comparison. (D) Representative flow cytometry profiles for landing pad cells expressing Parkin wild-type (n = 9.5×10^3^) and R42P (n = 1×10^4^) treated with 1 µM TAK243 for 6 hours. The untreated cells (from panel C) are included for comparison.

### Genome-wide CRISPR screening for components regulating PQC

To identify components involved in the PQC of R42P, we applied the Toronto CRISPR knockout library (TKOv3) (38) to HEK293T cells expressing the GFP-Parkin R42P variant. This library contains 3-4 sgRNAs targeting each of 18,053 different human protein-coding genes for a total of 70,948 guides. The sgRNAs were introduced in the HEK293T cells together with Cas9 by lentiviral transduction (**Fig. 2A**). The cells were then sorted by fluorescence-activated cell sorting (FACS) to isolate the 5% cells with the highest and lowest GFP:mCherry ratios (**Fig. 2A**), followed by sequencing across the sgRNAs to identify the genes that, when disrupted, lead to significant changes in R42P abundance (**Fig. 2A**). The screen revealed that the disruption of 74 genes resulted in significantly increased GFP-R42P:mCherry levels. Among these, we noted that the best scoring hits were the chaperone BAG6 and its associated E3 ubiquitin ligase RNF126 (**Fig. 2B**). Previous studies have shown that BAG6 forms a chaperone complex with UBL4A and GET4 (also known as TRC35) which, along with the co-chaperone SGTA, are involved in PQC (39). Accordingly, GET4 also scored as a hit in our screen (rank 14, p-value 2.39×10^-5^). UBL4A did not score using a p-value cut-off of 10^-3^ but would have been significant with a lower stringency (rank 193, p-value 4×10^-3^). SGTA was not a significant hit in our screen. As expected, the control sgRNAs targeting EGFP were the best scoring hit in the group displaying the 5% lowest GFP:mCherry ratios (**Fig. 2C**). In addition, this group contained 96 significant hits, including HSP90α (HSP90AA1) and HSP90β (HSP90AB1), the HSP90 co-chaperone STIP1 and the HSP70 co-chaperone DNAJA2 (**Fig. 2C**). We note that HSP90 scoring in the low 5% group is consistent with our results using geldanamycin (**Supplementary Fig. 1**), and also in agreement with the HSP90 inhibitory co-chaperones TSC1 and TSC2 (40) scoring as significant hits (ranked 4 and 6, respectively) in the high abundance pool (**Fig. 2B**). The sgRNAs to Parkin (*PRKN*/*PARK2*) did not score significantly, presumably due to Parkin R42P being codon optimized. Since the cells were cultured both before and after sorting, we do not expect essential PQC components, such as E1 and proteasome subunits, to be well represented in the screen. All screening results are included in the supplementary material (**Supplementary File 1**).

**Fig. 2.**
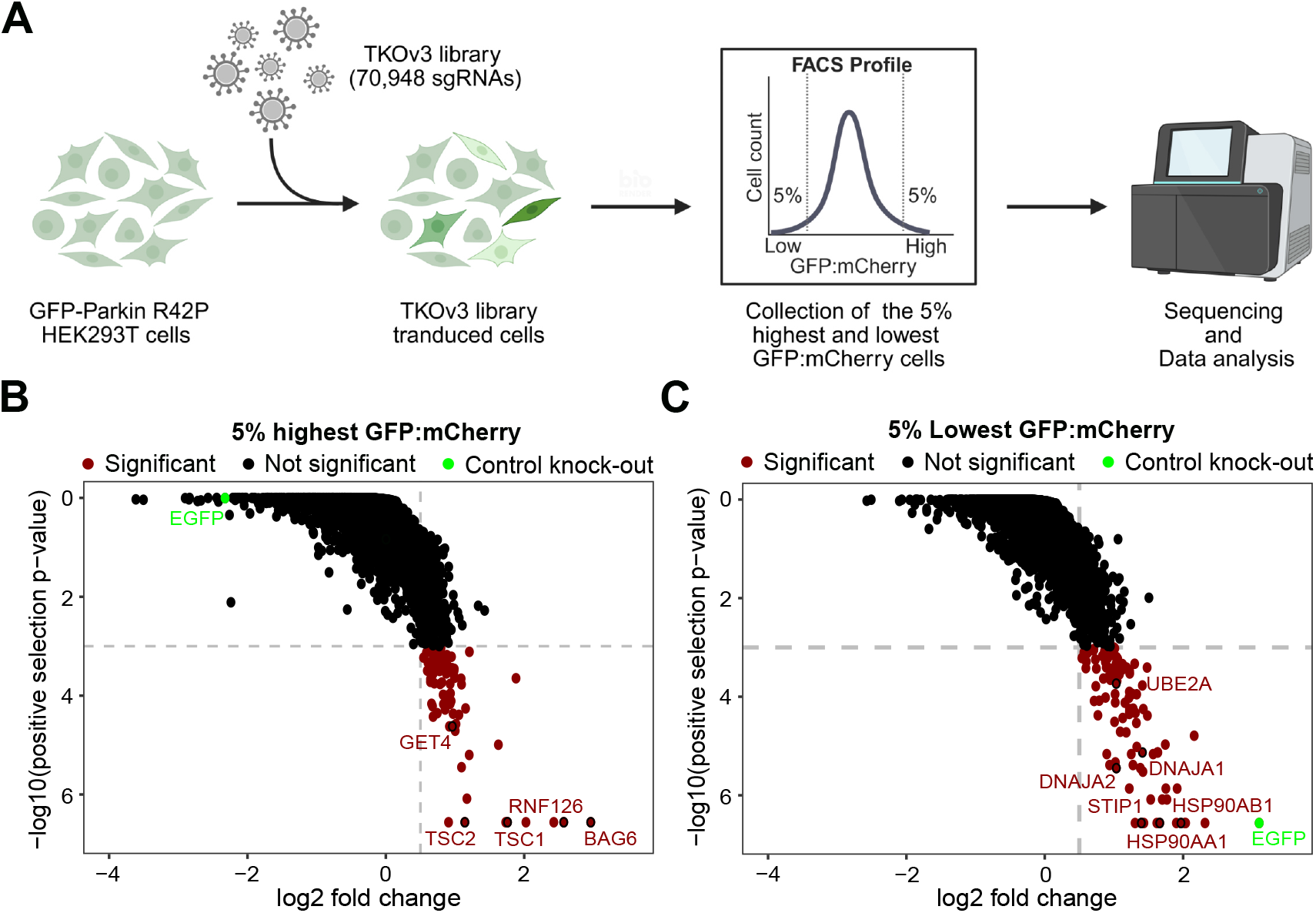
Genome-wide CRISPR screening for modulators of Parkin R42P abundance. (A) Schematic representation of the screening protocol. HEK293T cells expressing GFP-Parkin R42P from the landing pad were transduced with the TKOv3 sgRNA library. Fluorescence activated cells sorting (FACS) was used to collect the cells with the 5% highest and lowest GFP:mCherry ratios. Finally, the sgRNAs were identified by sequencing. Created in BioRender. Hartmann-Petersen, R. (2025) https://BioRender.com/mxx8owh. (B) Plot displaying sgRNA enrichment in the 5% highest GFP:mCherry bin. Significant (p < 0.001 and log2 fold change > 0.5) hits are colored red. Control sgRNAs are marked in green. (C) Plot displaying sgRNA enrichment in the 5% lowest GFP:mCherry bin. Significant (p < 0.001 and log2 fold change > 0.5) hits are colored red. Control sgRNAs are marked in green.

### BAG6 and RNF126 regulate PQC of Parkin R42P

To assess the validity of BAG6 and RNF126 scoring in the high 5% group, we generated their CRISPR knockouts in the HEK293T landing pad cell line. Although SGTA was not a significant hit in the screen, we also generated SGTA knockout cells due to its known involvement with BAG6. Western blotting revealed successful gene disruption (**Supplementary Fig. 2**). In addition, we noted that in BAG6 knockout cells the levels of UBL4A and GET4 were strongly reduced (**Supplementary Fig. 2**). However, this was not the case in RNF126 and SGTA knockout cells (**Supplementary Fig. 2**), indicating that while BAG6 is of key importance for the stability of the chaperone complex, SGTA and RNF126 are non-obligate BAG6 partners that are stable independently of BAG6. In accordance with the results from the screen, introducing wild-type Parkin and the R42P variant into the BAG6 and RNF126 knockout cells revealed an increase in abundance of the R42P variant (**Fig. 3A,B**). Re-introducing BAG6 and RNF126 into their respective knockout cells by transient transfection reduced the abundance of R42P (**Supplementary Fig. 3**). The abundance of R42P appeared largely unchanged in the SGTA knockout cells (**Supplementary Fig. 4**). In the BAG6 and RNF126 knockout cells, addition of bortezomib (**Fig. 3C,D**) or the E1 inhibitor TAK243 (**Fig. 3E,F**) led to a further increase in R42P abundance, suggesting that while BAG6 and RNF126 appear as key factors for Parkin R42P degradation, other UPS components also contribute to degradation.

**Fig. 3.**
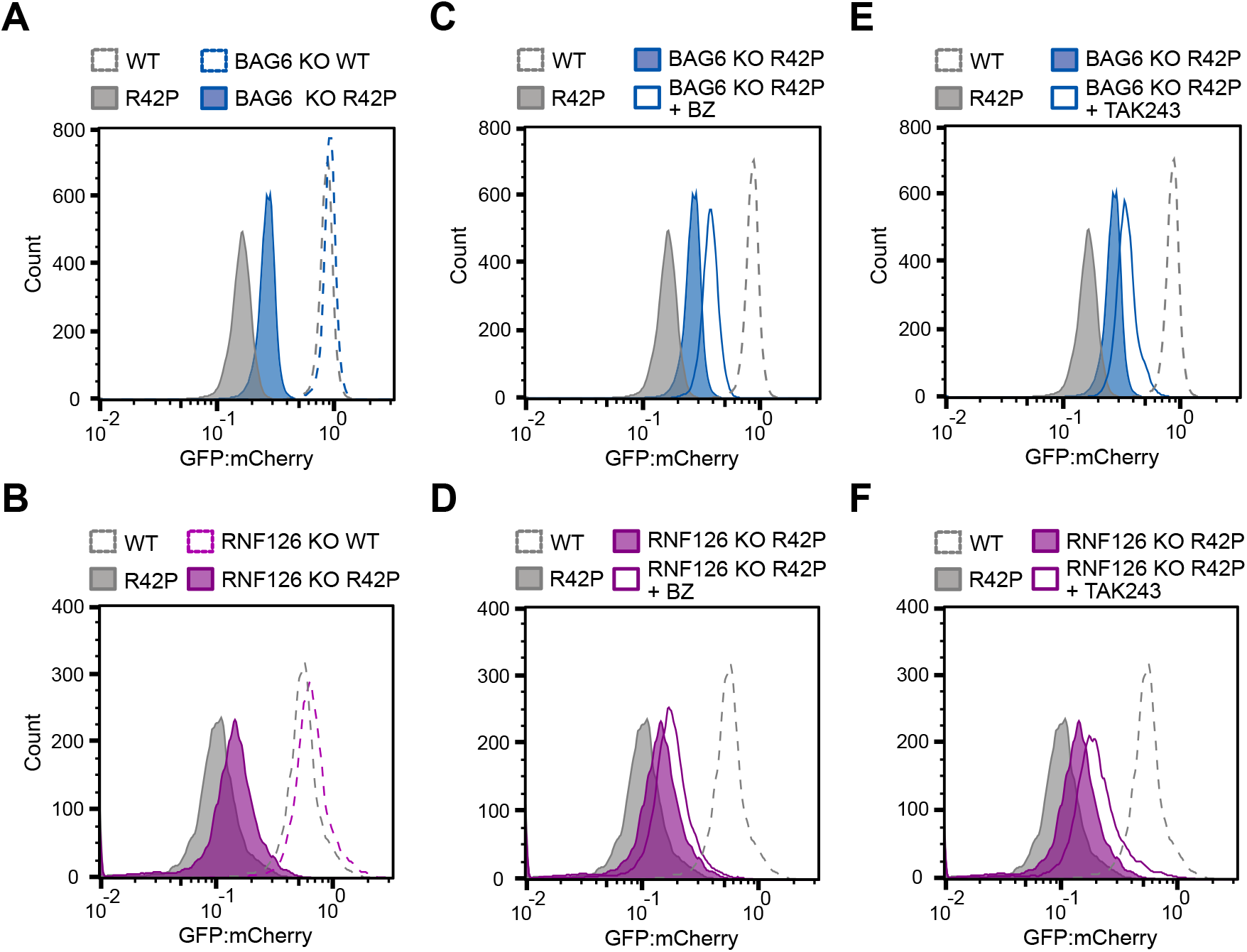
BAG6 and RNF126 regulate Parkin R42P abundance. Comparison of the steady-state protein levels of wild-type Parkin and R42P in (A) wild-type cells (WT n = 9.7×10^3^, R42P n = 1×10^4^) and BAG6 knockout cells (WT n = 1×10^4^, R42P n = 1×10^4^) and (B) wild-type cells (WT n = 9×10^3^, R42P n = 9×10^3^) and RNF126 knockout cells (WT n = 9×10^3^, R42P n = 9×10^3^). (C) Treatment with 10 µM bortezomib (BZ) for 16 hours in BAG6 knockout cells (R42P n = 9.5×10^3^) and (D) RNF126 knockout cells (R42P n = 9×10^3^). Controls from panels (A) and (B) are included for comparison. (E) Treatment with 1 µM TAK243 for 16 hours in BAG6 knockout cells (R42P n = 1.1×10^4^) and (F) RNF126 knockout cells (R42P n = 9×10^3^). Controls from panels (A) and (B) are included for comparison.

To test if Parkin and BAG6 interact, GFP-tagged wild-type Parkin, R42P, and as a control, GFP, were immunoprecipitated from HEK293T cells and analyzed by western blotting. The results showed that BAG6 was associated with R42P, but surprisingly also with wild-type Parkin (**Supplementary Fig. 5**). However, considering that the abundance of R42P is reduced (**Supplementary Fig. 5**), it seems that BAG6 may preferentially bind R42P. In conclusion, our results show that BAG6 and RNF126 selectively target Parkin R42P for degradation.

### Most low abundance Parkin missense protein variants are BAG6 and RNF126 targets

To further analyze the role of BAG6 and RNF126 in the PQC of Parkin missense variants, we utilized our previously described site-saturated and barcoded library of Parkin variants (31). This cDNA library contains 8757 out of 8835 (465 residues × 19 amino acid substitutions per position) possible single amino acid substitution variants and 462 of 464 (465-1 positions for early stop codons) nonsense variants, corresponding to >99% coverage. The Parkin variant library was introduced into the landing pad in wild-type HEK293T cells, as well as in the BAG6 and RNF126 knockout cell lines. The abundance of the Parkin variants in these different genetic backgrounds was then compared by flow cytometry. Consistent with our previous observations (31), the Parkin variant library displays a bimodal distribution, with a large peak representing variants with a wild-type abundance, and a smaller population of low abundance variants (**Fig. 4**). For both the BAG6 (**Fig. 4A**) and RNF126 (**Fig. 4B**) knockout cells, the shoulder of low abundance variants shifted towards higher GFP:mCherry ratios, indicating that most low abundance Parkin variants are targeted by BAG6 and RNF126. In agreement with our results on the R42P variant, SGTA knockout did not stabilize the low abundance Parkin variants in the library (**Supplementary Fig. 6**).

**Fig. 4.**
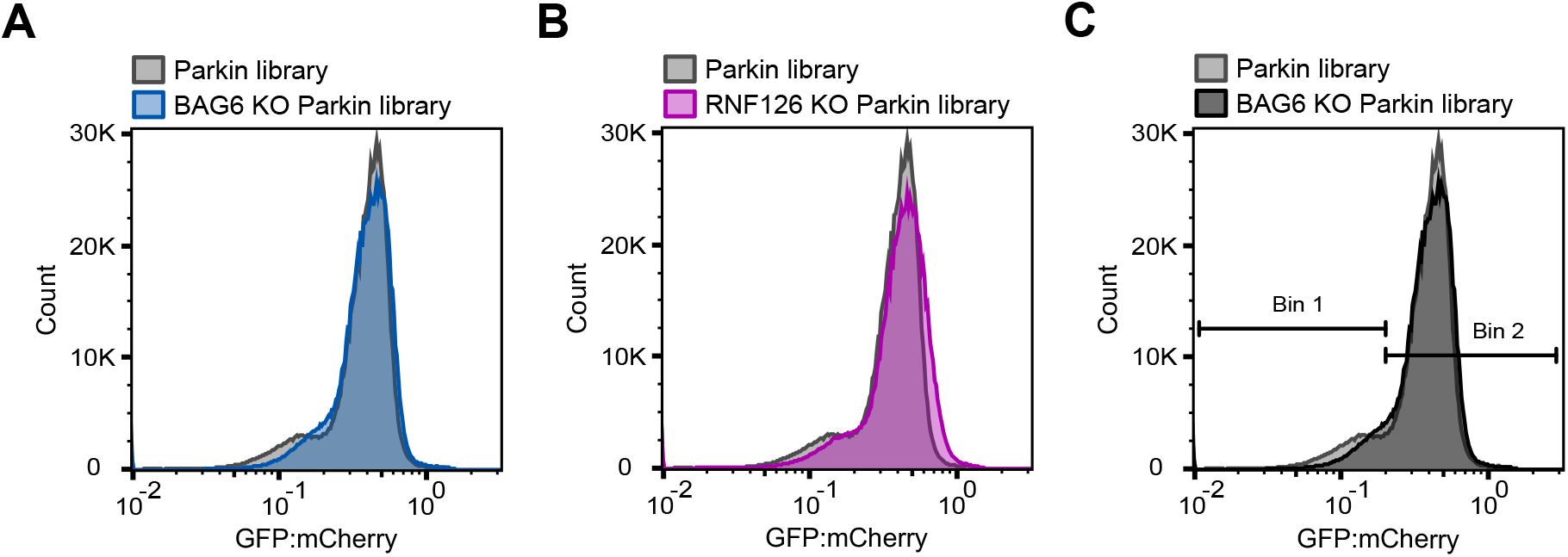
Most low abundance Parkin variants are BAG6 and RNF126 targets. Comparison of the flow cytometry profiles of the site saturated Parkin variant library expressed in wild-type cells (n = 1×10^6^) and (A) BAG6 knockout cells (n = 1×10^6^) and (B) RNF126 knockout cells (n = 1×10^6^). (C) As panel (A) including the gating scheme used for flow sorting.

To pinpoint those Parkin variants that are targeted by BAG6, we used the variant abundance by massively parallel sequencing (VAMP-seq) method (28) where cells are sorted into bins and Parkin variants identified by sequencing. The library was expressed in both wild-type cells and BAG6 knockout cells to determine abundance changes under the same experimental settings. Focusing on low abundance variants, we sorted cells into two categories using fixed bins derived from the GFP:mCherry distribution in wild-type cells. This corresponded to 17% and 5% of cells assigned to bin 1 in wild-type and BAG6 knockout cells, respectively, with the remaining cells assigned to bin 2 (**Fig. 4C**). Sequencing the barcodes allowed us to determine a protein stability index (PSI) for each variant in wild-type and BAG6 knockout cells (see Methods). The difference in the protein stability index (ΔPSI) between wild-type and BAG6 knockout cells revealed the BAG6-responsive variants (high ΔPSI). The R42P variant displayed a ΔPSI of 0.38 which we use as a threshold for identifying BAG6 targets. We identified 1518 variants with low abundance in the wild-type control cells (PSI < 1.5), of which 1128 variants (74%) displayed an increased abundance (ΔPSI ≥ 0.38) in the BAG6 knockout cells. Comparison with the previously determined Parkin variant abundance (31) (**Fig. 5A**) revealed that these BAG6-targets (increased abundance) variants were distributed throughout the structured regions in Parkin (**Fig. 5B**). Surprisingly, only a few nonsense variants were included in this group, perhaps because they were too low in abundance in both wild-type and BAG6 knockout cells and therefore poorly resolved (i.e. falling below the FACS gating threshold and therefore did not switch bins upon BAG6 disruption). Alternatively, some early stop codons could generate C-degrons that are BAG6 independent (12, 41, 42). Of the 390 non-targets (low abundance with ΔPSI < 0.38), 328 (84%) were nonsense variants and the remaining 62 were primarily located towards the N- and C-terminal regions of the protein (**Supplementary Fig. 7**). As expected, the low abundance BAG6 targets identified here were also measured to be of low abundance in the previous VAMP-seq (**Fig. 5C,D**) and primarily found at positions with a low relative accessible surface area (rASA) (**Supplementary Fig. 8A**) and predicted (by Rosetta) to result in significant structural destabilization (**Supplementary Fig. 8B**). In agreement with this, mapping the median (mdn) ΔPSI per position onto the Parkin structure, revealed that most BAG6 responsive variant positions were buried in the structure (**Fig. 5E**).

**Fig. 5.**
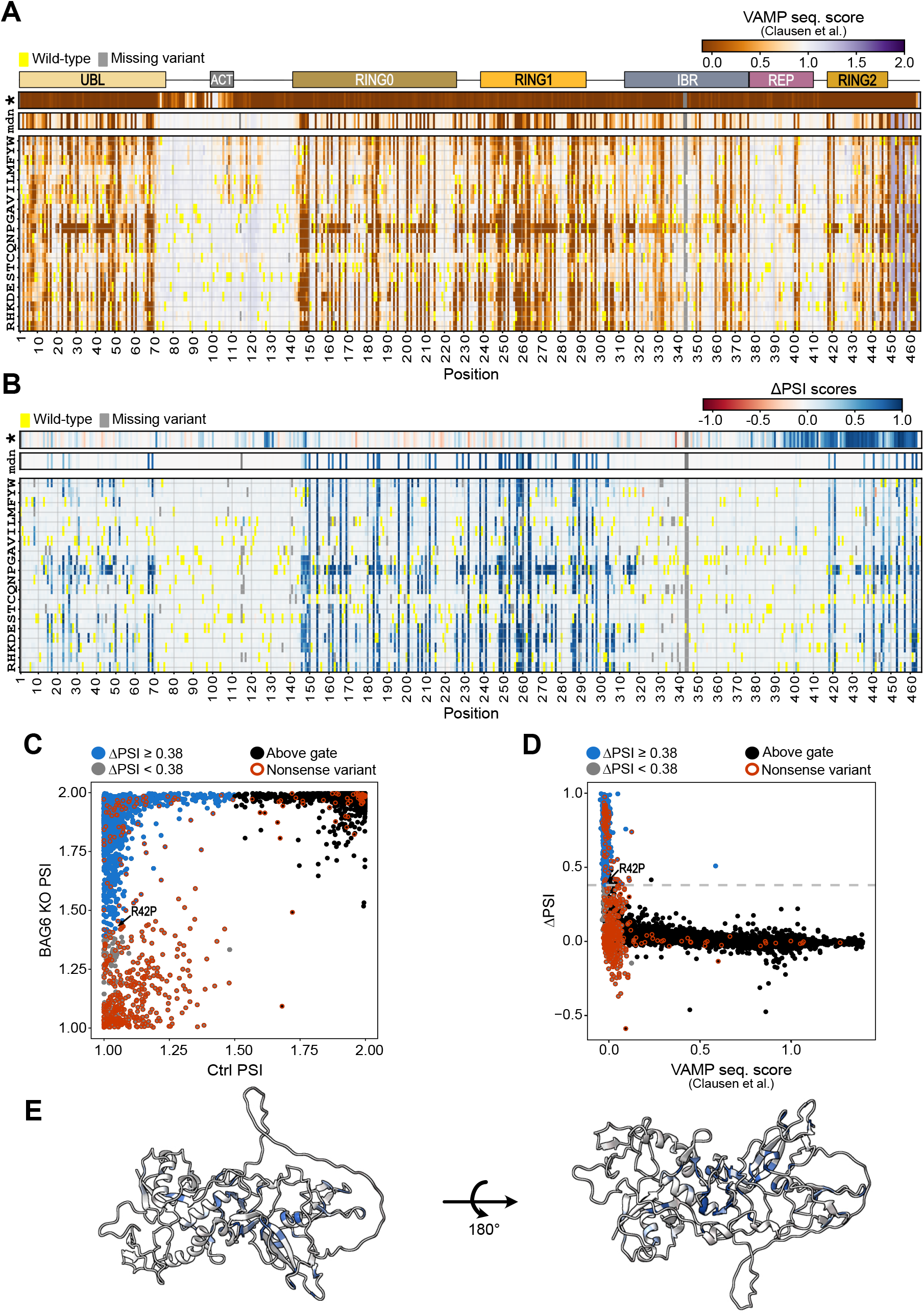
Mapping the low abundance Parkin targets of BAG6. (A) Heatmap of the Parkin variant abundance scores determined previously by VAMP-seq (31). The nonsense variants, marked with an asterisk (*) and median abundance scores (mdn) per position for missense variants are shown above the heat-map. The abundance scores range from low (red) over WT-like (white) to increased abundance (blue). Missing variants are marked in grey, and the wild-type residues are shown in yellow. The Parkin domain organization is indicated with colored bars. (B) Heatmap showing the Parkin variants abundance change in BAG6 knockout cells. The colors display the ΔPSI, with blue variants being stabilized. (C) Comparison of the PSIs in wild-type (Crtl) cells and BAG6 knockout cells. Low abundance variants were defined as variants with a control PSI < 1.5 and colored blue or grey according to ΔPSI. High abundance variants (control PSI ≥ 1.5), termed above gate are marked in black (above the FACS gate). (D) Comparison of the Parkin variant abundance in wild-type cells as determined before (31) (x-axis) and the ΔPSIs in the BAG6 knockout cells determined here. The new VAMP-seq experiment was designed with focus on low abundance variants, while the previous data also resolved the high abundance variants. (E) The AlphaFold2 predicted full-length Parkin structure (AF-O60260-F1) colored by the median (mdn) ΔPSI as in panel B. Note that the BAG6-responsive positions (blue) are buried.

Finally, we assessed ΔPSIs of the low abundance pathogenic missense variants that we identified before (31). In all cases, these scored as BAG6 targets (ΔPSI ≥ 0.38) (**Supplementary Table 1**). We conclude that BAG6 disruption broadly increases the abundance level of non-native Parkin variants, including some pathogenic variants.

### BAG6 and RNF126 are broadly involved in PQC of missense protein variants

Having established that many structurally destabilized Parkin variants are targeted by BAG6 and RNF126, we continued to test if this was unique for Parkin or if other destabilized missense protein variants were also targets. To this end, we selected two other model proteins, the tumor suppressor folliculin (FLCN), linked to Birt-Hogg-Dubé syndrome (OMIM: 135150) (43), and phenylalanine hydroxylase (PAH) linked to phenylketonuria (OMIM: 261600) (44). Previously, we have shown that the pathogenic FLCN H255P and R239C, and PAH I65T and L348V variants are destabilized PQC targets (45, 46). Indeed, introducing these variants into the wild-type HEK293T landing pad showed a reduced abundance compared to the wild-type proteins (**Fig. 6A,B**). For FLCN H255P, FLCN R239C, and PAH L348V, the abundance was increased in the BAG6 and RNF126 knockout cells, while the level of PAH I65T was unchanged (**Fig. 6A,B**). Based on these data, we conclude that BAG6 and RNF126 are broadly involved in PQC of missense protein variants.

**Fig. 6.**
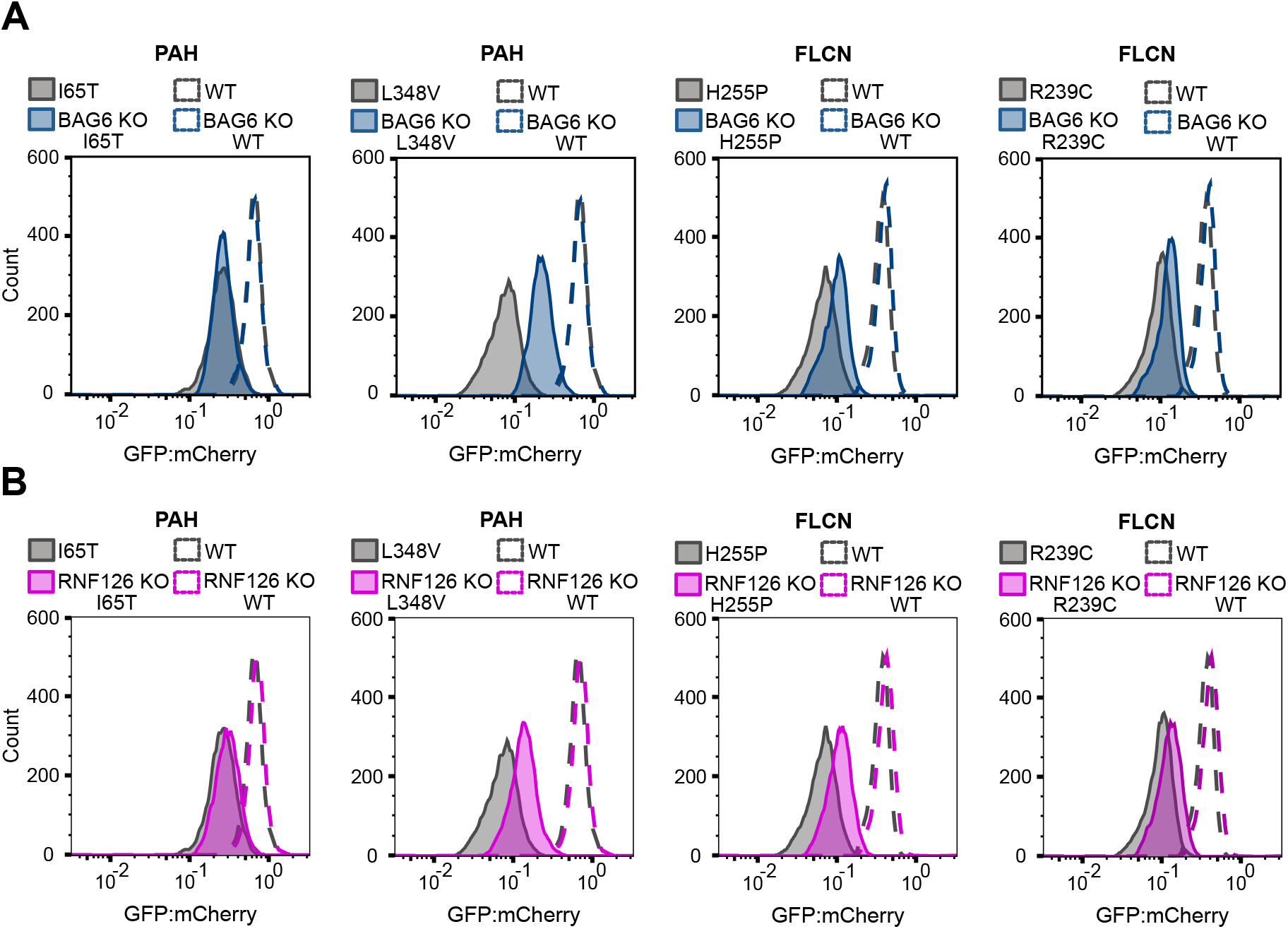
Low abundance PAH and FLCN variants are BAG6 and RNF126 targets. (A) Comparison of the steady-state protein levels of wild-type (WT) PAH, PAH I65T, PAH L348V, FLCN, FLCN H255P and R239C variants in wild-type and BAG6 knockout cells. (B) Comparison of the steady-state protein levels of wild-type (WT) PAH, PAH I65T, PAH L348V, FLCN, FLCN H255P and R239C variants in wild-type and RNF126 knockout cells. N = 10^4^ cells for all experiments.

## Discussion

The results presented here show that the chaperone BAG6 along with its associated E3, RNF126, contribute to the PQC of most low abundance and non-native Parkin variants, including several pathogenic variants. Additionally, structurally destabilized variants of the unrelated proteins FLCN and PAH were also BAG6 and RNF126 targets, indicating that BAG6 and RNF126 are likely involved more broadly in PQC of missense variants. These results are in accordance with recent CRISPR screens that have established a connection between PQC degrons and BAG6 (13). Importantly, the same study also revealed that the BAG6-dependent degrons can be as short as five consecutive hydrophobic residues. The seeming promiscuity for short hydrophobic stretches (13) is compatible with the BAG6 complex regulating PQC turnover of missense protein variants, and we note that several PQC degrons are embedded throughout the Parkin sequence (31). Since the regions flanking I65 in PAH are rich in hydrophilic and charged residues that are not compatible with a PQC degron, this could also potentially explain why a locally unfolded PAH I65T variant is not a target of BAG6 and RNF126.

Previous studies have shown that BAG6 forms an ATP-independent chaperone complex with UBL4A and GET4, that together with the SGTA co-chaperone facilitates PQC of tail-anchored proteins and mislocalized transmembrane proteins (19, 20, 22-24, 39, 47-49). When membrane insertion fails, RNF126 catalyzes substrate ubiquitylation leading to proteasomal degradation of the non-native target protein (19, 22, 23, 48, 49). The results presented here show that BAG6 and RNF126 are also critical for degradation of structurally destabilized missense protein variants and thus appear broadly connected to cytosolic PQC. Since GET4 also scored in our screen and UBL4A would have scored with a 1% significance threshold, it is likely that the entire BAG6:GET4:UBL4A complex is involved in triage of missense protein variants. On the other hand, UBL4A and GET4 might primarily regulate substrate transfer and membrane insertion (39), potentially explaining their lower scores compared to BAG6. In addition, BAG6 stability might not depend on UBL4A and GET4, while we conversely show that BAG6 is essential for UBL4A and GET4 stability. The SGTA co-chaperone did not score in our screen, and if anything, the Parkin variants appeared slightly less abundant in the SGTA knockout cells, which may support an antagonistic role of SGTA in BAG6-mediated triage, as earlier noted (50). Previous data also indicate that BAG6 may directly recognize targets (20, 23, 24, 51), so triage by the BAG6 complex could occur independently of SGTA. Although a recent study has provided structural information on the BAG6 complex (52), this does not include the ∼900 residue region in BAG6 that is likely involved in substrate binding. Thus, details on substrate binding and selection await further structural characterization. The GET3 (also known as ASNA1 or TRC40) co-factor of the BAG6 complex, which has been associated with membrane insertion of tail-anchored proteins downstream of SGTA and BAG6 (53-55), did not score in our screens. This may suggest that the BAG6 complex selectively routes destabilized missense protein variants for degradation, but it is unknown how the fate of BAG6 substrates is determined.

We found that inhibiting E1 or the proteasome in BAG6 and RNF126 knockout cells leads to further stabilization of Parkin R42P. This indicates that at least one other E3 must work in parallel with BAG6 and RNF126. Potentially, this could include UBE3C, which also scored in our screen (rank 35). However, multiple E3s are likely to contribute to PQC degradation. Thus, it appears that cells may rely on both the broad substrate selectivity of molecular chaperones and several E3s with overlapping substrate specificities. This is in accordance with the previously noted functional redundancy between PQC E3s (56-58). Although most components of the PQC and UPS systems are conserved in eukaryotes, this does not include some of the PQC E3s. For instance, efforts to identify a human ortholog of the yeast ubiquitin ligase San1 have failed (59), and conversely, the yeast genome does not encode an ortholog of CHIP. While RNF126 seems to be distantly related to fission yeast Bop1 (SPAP32A8.03c), BAG6 is not found in any yeast species. It therefore appears that the arsenal of PQC components and degradation pathways display some notable differences between species.

Our results also show that HSP90 likely assists in the folding of Parkin, so when HSP90 is inhibited or disrupted, a larger fraction of Parkin is unable to attain its native fold, leading to degradation and reduced protein levels. This is in agreement with HSP90 targeting some clients for degradation (60), and we note that the HSP90 co-chaperone STIP1 also scored as a hit in the low abundance pool. TSC1 and TSC2 have been shown to compete with the activating co-chaperones for binding to HSP90 (40), which is in line with TSC1/2 scoring in the high abundance pool. Our observation that deletion of DNAJA2 reduces the abundance of R42P suggests that DNAJA2 contributes to Parkin folding and stability. This agrees with the well-characterized role of DNAJA2 in protein folding (61) and that DNAJA2 buffers proteasomal degradation of unstable proteins (62).

In conclusion, our results support a model where structurally destabilized missense protein variants are folded or stabilized by HSP90 and STIP1, while BAG6 and RNF126 contribute to their PQC degradation (**Fig. 7**). The system appears broadly inclusive, suggesting that many missense protein variants follow this route. Interfering with this pathway may offer a strategy to boost the level of structurally destabilized, but functional, pathogenic protein variants to suppress the loss-of-function phenotypes and mitigate disease.

**Fig. 7.**
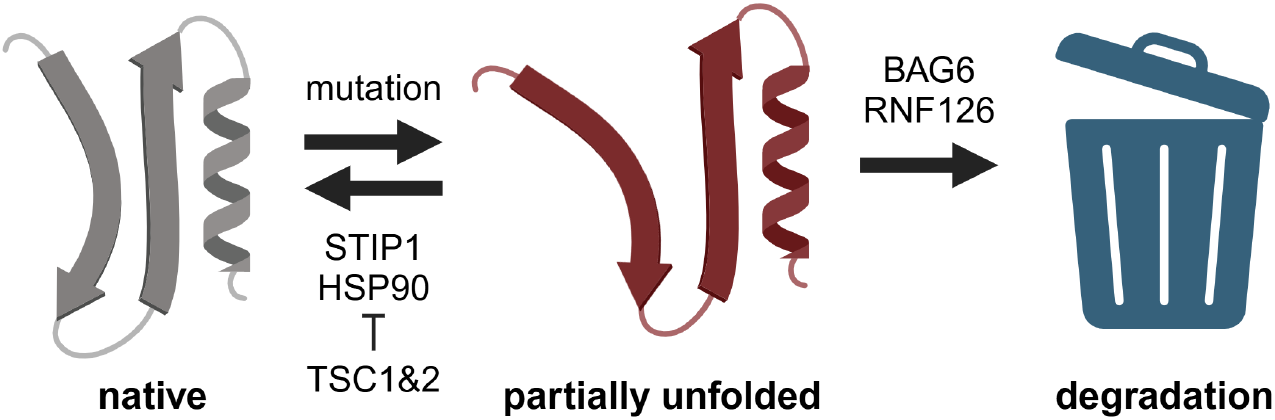
Model of the PQC and degradation of missense proteins. Mutations leading to a partial (or full) unfolding of the native structure render the protein a target for BAG6 and RNF126-mediated ubiquitin-dependent proteasomal degradation. Conversely, HSP90 and STIP1 stabilize the native state, while TSC1 and TSC2 inhibit HSP90. Created in BioRender. Hartmann-Petersen, R. (2025) https://BioRender.com/sly8ce1.

## Methods

### Plasmids

The full-length *PRKN* wild-type and R42P cDNA were codon optimized for expression in human cells and inserted into a VAMP-seq expression vector (28) with EGFP fused N-terminally (Genscript), as described before (31). The empty vector control was the VAMP-seq expression vector without cDNA fused to GFP. The recombinase Bxb1 was expressed from pCAG-NLS-HABxb1 (Addgene #51271) (63). The *PRKN* cDNA library has been described before (31). The Toronto human knockout pooled library (TKOv3) (Addgene #90294) (64), psPAX2 (Addgene #12260) and pMD2.G (Addgene #12259) were from Addgene. BFP and myc-BAG6-His were expressed transiently from pcDNA3.1 and RNF126-HA from pcDNA5/FRT/TO (Genscript).

### Cell culture and transfections

The HEK293T TetBxb1BPFiCasp9 Clone 12 cell line (37) was cultured in Dulbecco’s Modified Eagle High-Glucose medium (DMEM) (Sigma-Aldrich) supplemented with 10% (v/v) fetal bovine serum (FBS) (Sigma Aldrich), 5 mg/mL streptomycin (BioChemica), 5000 UI/mL penicillin (BioChemica), and 2 mM L-glutamine (Sigma Aldrich) at 37 °C in a humidified incubator containing 5% CO_2_. cDNA of single variants and the barcoded library were transfected and integrated into the Tet-on landing pad using the FuGENE HD transfection reagent (Promega) with a 4:1 ratio of transfection reagent to DNA and a 14:1 ratio of *PRKN* plasmid to Bxb1 plasmid. Gene expression from the Tet-on promoter was activated by 2 µg/mL doxycycline (Sigma Aldrich) and apoptosis was induced in non-recombinant cells by 10 nM of AP1903 (MedChemExpress) about 48 hours after transfection. Cells were kept for no longer than 18 passages and transfected cells were supplemented with 2 µg/mL doxycycline at each passage. Transient transfections were carried out five days after Tet-on promoter activation using the FuGENE HD transfection reagent (Promega) at a 4:1 ratio of transfection reagent to DNA. The DNA was a mixture of BAG6 or RNF126 plasmid and BFP plasmid at a 10:1 ratio.

### CRISPR screening

The Toronto Knockout v3 (TKOv3) CRISPR library in lentiCRISPR v2 backbone (LCV2) (Addgene #90294) (64) was used in the pooled CRISPR-Cas9 knockout screen of Parkin R42P abundance. TKOv3 CRISPR library amplification, large-scale lentivirus production of TKOv3, cell line characterization, and lentivirus multiplicity of infection (MOI) determination were performed according to manufacturer’s protocol (Addgene) with the following deviations. During lentivirus production lipofectamine3000 and P300 (Thermo Fisher Scientific) were used in place of XtremeGENE 9. Transfected cells were incubated for 6 hours in a 37 °C humidified incubator containing 5% CO_2_. After 6 hours, the medium containing transfection mixture was replaced with pre-warmed DMEM supplemented with 10% (v/v) FBS, 1% bovine serum albumin (BSA), 5 mg/mL streptomycin, and 5000 UI/mL penicillin, then returned to the incubator for an additional 48 hours of incubation. The supernatant was collected by centrifugation at 300 g for 5 minutes and filtered through a 0.45 µm filter before being aliquoted and stored at -80 °C.

The primary screen was performed in the HEK293T TetBxb1BPFiCasp9 Clone 12 cells expressing the Parkin R42P variant. R42P transfected cells were expanded to at least 8.3×10^7^ cells to ensure a minimum coverage of 350-fold of the CRISPR sgRNA library (71,090 sgRNA) with an MOI at ∼0.3. The R42P transfected cells were pooled and infected with TKOv3 lentivirus supplemented with 8 µg/mL polybrene (Sigma-Aldrich) to increase infection efficiency. The transduced cells were seeded at 30 % confluency in 15 cm tissue culture dishes and incubated for 24 hours in a 37 °C humidified incubator containing 5% CO_2_. After 24 hours, the transduced cells were treated for 48 hours with 2 µg/mL puromycin (Thermo Fisher Scientific) to select the infected cells. The MOI was determined by dividing the cell concentration of a plate after puromycin treatment with the cell concentration of a plate not treated with puromycin. The MOI of the infected cells was 0.38 and the coverage was 900-fold. In total 5×10^7^ cells were collected as timepoint zero (T0) by centrifugation at 300 g for 7 minutes and the cell pellet was stored at -80 °C. The remaining infected cells were seeded at 20% confluency. The cells were cultured in DMEM supplemented with 10% (v/v) FBS, 5 mg/mL streptomycin, 5000 UI/mL penicillin, 2 mM L-glutamine, and 2 µg/mL doxycycline and passaged every third day. The coverage was kept at 350-fold. After 12 days (from T0), cells were washed in PBS, dislodged with trypsin and resuspended through a 50 µm nylon mesh filter in PBS containing 2% (v/v) FBS. A sample of 5×10^7^ cells was collected prior to Fluorescence-Activated Cell Sorting (FACS) by centrifugation at 300 g for 7 minutes and the cell pellet was stored at -80 °C. FACS was performed using a BD FACSymphony S6 cell sorter (BD Biosciences). Blue fluorescent protein (BFP) expressed in non-recombinant cells was excited with a 405 nm laser and emitted light was collected after passing through a 410 long pass filter and a 431/28 nm band pass filter. EGFP expressed from the recombined landing pad was excited with a 488 nm laser, and emitted light was collected after passing through a 505 nm long pass filter and a 530/30 nm band pass filter. mCherry was excited with a 561 nm laser, and emitted light was collected through a 600 nm long pass filter followed by a 610/20 nm band pass filter. For sorting, the cells were gated as live, singlets, BFP negative, mCherry positive, and sorted based on the GFP:mCherry ratio. The gating strategy is illustrated in the supplementary material (**Supplementary Fig. 9**). A total of 3.5×10^6^ cells with the 5 % highest and lowest GFP:mCherry ratio were collected (50-fold coverage). The collected cells were cultured without doxycycline for 5 days, before 5×10^7^ cells were harvested and stored -80 °C.

The genomic DNA was extracted from the cells as described (65), except that after air drying the DNA was resuspended in 300 µL of 0.1x TE buffer instead of 500 µL of 1x TE buffer (10 mM Tris/HCl, 1 mM EDTA, pH 8).

The TKOv3 CRISPR sequencing library preparation was as described in the Addgene protocol with few deviations. PCR 1 was executed with 20 cycles instead of 25 cycles. The PCR 2 product was re-purified with GeneJET PCR Purification Kit (Thermo Fisher Scientific) after gel extraction. Primers are listed in supplementary material (**Supplementary Table 2**).

The knockout libraries were pooled, denatured and mixed with PhiX Control v3 following manufacturer’s protocol (Illumina). The prepared libraries were sequenced using a NextSeq 550 sequencer with a NextSeq 500/550 High Output v2.5 75 cycle kit (Illumina) with standard primers.

### Single CRISPR knockout cell lines

The BAG6 CRISPR/Cas9 knockout plasmid (sc-432477, Santa Cruz), BAG6 HDR plasmid (sc-432477-HDR, Santa Cruz Biotechnology), and Cre Vector (sc-418923, Santa Cruz Biotechnology) were used to generate HEK293T TetBxb1BPFiCasp9 BAG6 knockout cells. First, 0.5 µg BAG6 CRISPR/Cas9 knockout plasmid, 0.5 µg BAG6 HDR plasmid, 100 µL OptiMEM (Thermo Fisher Scientific) and 10 µL FuGENE HD transfection reagent (Promega) were mixed and incubated at RT for 15 minutes. The transfection mixture was added dropwise to cultured cells and incubated for 48 hours in a 37 °C humidified incubator containing 5% CO_2_. To select for stable transfected cells and activate the landing pad, 1 µg/mL puromycin (Thermo Fisher Scientific) and 2 µg/mL doxycycline (Sigma Aldrich) were used. After selection, 1 µg Cre Vector plasmid, 100 µL OptiMEM, and 10 µL FuGENE HD transfection reagent were mixed and incubated 15 minutes at RT, then added to the transfected cells to remove the red fluorescent protein (RFP) from the integrated HDR plasmid. After 48 hours incubation, the cells were diluted to isolate single cells in a 96-well plate. Successful knockout was confirmed by western blotting.

For RNF126 (sc-405024, Santa Cruz Biotechnology) and SGTA (sc-404399, Santa Cruz Biotechnology) knockouts, 1 µg of the CRISPR/Cas9 knockout plasmid, 100 µL OptiMEM, and 10 µL FuGENE HD transfection reagent were mixed and incubated for 15 minutes at RT. The transfection mixture was added dropwise to cultured cells and incubated for 24-28 hours in a 37 °C humidified incubator containing 5% CO_2_. After incubation, the cells were washed in PBS, detached with trypsin and resuspended through a 35 µm nylon mesh filter in PBS containing 2% FBS. To select stable transfected cells and isolate single colonies, the cells were sorted based on GFP levels from the transient plasmid on a BD FACSJazz cell sorter (BD Biosciences). GFP was excited with a 488 nm laser, and emitted light was collected after passing through a 530/40 nm band pass filter. In addition, the cells were gated for live singlets using FSC and SSC (for live cells), FSC and Trigger Pulse Width (for singlets). The cells were single cell seeded into 96-well plates containing 50% conditioned medium in DMEM supplemented with 10% (v/v) FBS, 5 mg/mL streptomycin, 5000 UI/mL penicillin, 2 mM L-glutamine, and 2 µg/mL doxycycline. Successful knockout was confirmed by western blotting.

### Immunoprecipitation

Cells were grown and transfected as described above. The cells were washed in PBS and harvested in IP lysis buffer (50 mM Tris/HCl pH 7.4, 150 mM NaCl, 1 mM EDTA, 0.5% NP-40, 1 mM PMSF, and 1 complete protease inhibitor (Roche)). The cell lysate was incubated at 4 °C for 30 minutes and centrifuged at 15,000 g for 20 minutes at 4 °C. A small amount of supernatant was mixed with SDS sample buffer (2% SDS, 62.5 mM Tris/HCl pH 6.8, 10% (v/v) glycerol, 0.005% bromophenol blue, 0.005% pyronin G, 1% (v/v) 2-mercaptoethanol) and used as input samples, whereas the remaining supernatant was mixed with 20 µL (bed volume) GFP-trap beads (Chromotek). Samples were tumbled for 4 hours at 4 °C followed by five wash steps by centrifugation in IP lysis buffer. The proteins were eluted in SDS sample buffer and boiled for 2 minutes. The samples were analyzed with SDS-PAGE and western blotting.

### SDS-PAGE and western blotting

Whole cell lysates were prepared by lysing harvested cells directly with SDS sample buffer. The whole-cell lysates were sonicated and boiled for 2 minutes. The proteins were resolved on 12.5% acrylamide separation gels with 3% stacking gels at a voltage of 125-150 V for 45-60 minutes. The gels were transferred to 0.2 µm nitrocellulose membranes (Adventec) at 100 mAmp/gel for 1.5 hours, followed by blocking of the membranes for at least 30 minutes in 5% (w/v) skim milk powder, 0.1% (v/v) Tween-20 and 2.5 mM NaN_3_ in PBS. Primary antibodies were applied to the membranes overnight at 4 °C, then washed extensively prior to at least 1 hour incubation in secondary antibody. The blots were developed using ECL detection reagents (GE Healthcare) and images were captured on a ChemiDoc MP Imaging System (BioRad). The antibodies were: anti-GFP (diluted 1:1,000) (Chromotek, 3H9), anti-mCherry (diluted 1:1,000) (Chromotek, 6G6), anti-β-actin (diluted 1:20,000) (Sigma, A5441), anti-BAG6 (1:5,000) (Proteintech, 66661-1-Ig), anti-RNF126 (1:5,000) (Proteintech, 66647-1-Ig), anti-SGTA (1:5,000) (Proteintech, 60305-1-Ig), HRP anti-mouse (diluted 1:5,000) (Dako, P0260), and HRP anti-rat (diluted 1:5,000) (Invitrogen, 31470).

### Flow cytometry

Flow cytometry was performed on transfected cells 6 days after the Tet-on promoter was activated. In the case of transient transfections, cells were analyzed 2 days after co-transfection. Perturbations with 10 µM bortezomib (LC Laboratories), 1 µM TAK-243 (MedChemExpress), and 1 µM geldanamycin (Sigma Aldrich) were performed 16 hours prior to flow cytometry. Cells were washed in PBS, detached with trypsin and resuspended through a 50 µm nylon mesh filter in PBS containing 2% FBS. The cells were analyzed using a BD FACSJazz cell sorter (BD Biosciences) with the following lasers 405 nm, 488 nm, 561 nm and band pass filters 450/50 nm, 530/40 nm and 610/20 nm for BFP, EGFP and mCherry, respectively. The cells were gated for live, singlets, BFP negative and mCherry positive cells using FSC-A and SSC-A (for live cells), FSC-A and Trigger Pulse Width (for singlets) with the BD FACS Software v.1.2.0.142. For transiently transfected cells, BFP and mCherry positive cells were gated, indicating the successful co-transfection of both plasmids. Back gating was performed to gate for the main population. Between 10^4^ and 10^6^ cells were recorded. Further analysis of the flow profiles was performed using FlowJo v.10.10.0 software (BD Biosciences). The full gating strategy is provided in the supplementary material (**Supplementary Fig. 10 and Supplementary Fig. 11**).

### VAMP-seq in the BAG6 knockout cells

The *PRKN* variant library was transfected into the HEK293T TetBxb1BPFiCasp9 BAG6 knockout Clone 1 cell line and the parental HEK293T TetBxb1BPFiCasp9 Clone 12 cell line as described previously (31). Then, 5 days after landing pad activation by 2 µg/mL doxycycline, cells were washed in PBS, detached with trypsin and resuspended through a 50 µm nylon mesh filter in PBS containing 2% FBS. FACS was performed using a BD FACSymphony S6 (BD Biosciences) cell sorter with the same laser and filters as described above. The cells were gated for live, singlets, BFP negative, mCherry positive, and the GFP:mCherry ratio. The cells were collected in two bins that were set based on the cell distribution profile of the GFP:mCherry ratio in the parental cell line and kept unchanged for the library in the BAG6 knockout cell line (**Supplementary Fig. 12**). The collected cells were seeded and grown to 10^7^ cells in doxycycline free medium before harvesting and storage at -80 °C.

Genomic DNA was extracted from the cells using the DNeasy blood & tissue kit (Qiagen). The genomic DNA was amplified from each sample with four 50 µL PCR reactions. Each reaction was prepared with 2 µg genomic DNA, 0.5 µM forward primer (LC1020), 0.5 µM reverse primer (LC1031) and 25 µL 2x Q5 High-Fidelity Mastermix (New England BioLabs). The initial denaturation was performed at 98□°C for 30□seconds, then 7 cycles of denaturation at 98□°C for 10□seconds, annealing at 60□°C for 20□seconds and extension at 72□°C for 10□seconds and a final extension at 72□°C for 2□minutes. The four reactions per sample were pooled and purified using AMPure XP beads (Beckman Coulter). The DNA was eluted in 16 µL nuclease-free water, and 7 µL of the DNA was used in a second PCR reaction to add the I5 and I7 Illumina cluster-generating sequences. Each reaction was prepared with 7 µL DNA, 0.5 µM I5 indexed primer (PCR2_Fw), 0.5 µM I7 indexed primer (JS_R), 10x SYBR green (Thermo Fisher Scientific) and 25 µL 2x Q5 High-Fidelity Mastermix. The initial denaturation was performed at 98□°C for 30□seconds, then 11 cycles of denaturation at 98□°C for 10□seconds, annealing at 63□°C for 20□seconds and extension at 72□°C for 15□seconds. The amplicons were loaded on a 2% agarose gel with 1x SYBR Safe (Thermo Fisher Scientific) and run for 75 minutes at 125 V followed by extraction using GeneJET gel extraction kit (Thermo Fisher Scientific). The samples were pooled, denatured and mixed with PhiX Control v3 following manufacturer’s protocol (Illumina). The prepared library was sequenced using a NextSeq 550 sequencer with a NextSeq 500/550 Mid Output v2.5 150 cycle kit (Illumina) with custom sequencing primers. Read 1 (LC1040), Read 2 (LC1041), Index 1 (LC1042) and Index 2 (ASPA_Park2 index2_Re). All primer sequences are listed in the supplementary material (**Supplementary Table 2**).

### Sequence data analyses

Sequencing reads were processed as in (31). Briefly, adapter sequences were removed using cutadapt (66) and paired-end reads were joined using ea-utils (67). Only exact matches to the sgRNA and barcode library were counted. Read counts from the CRISPR screen were processed using mageck version 0.5.9.5 (68).

Barcode read counts from the VAMP-seq were aggregated per amino acid variant and normalized to frequencies using a pseudo count of one. For each biological replica, a protein stability index (PSI) was calculated per variant using:

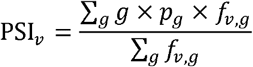

where *f*_*v,g*_ is the frequency of variant, *v*, in FACS gate *g*. Because we were comparing PSI values of two cell lines, the FACS gate was fixed in all sortings, and the population of cells in each gate, *p*_*g*_, was recorded to 0.17 and 0.83 for gate one and two for the control cells and 0.05 and 0.95 for the BAG6 knockout cells. We required 50 or more reads across both gates for a confident PSI, resulting in a coverage of 8738 of 8835 (99%) single variants and 462 of 464 (99%) nonsense variants measured in both cell lines. PSI values were well reproduced across the two biological replicates with Pearson correlations of 0.98 and 0.91 for control and BAG6 knockout cells, respectively.

## Abbreviations

BZ: bortezomib
FACS: fluorescence-activated cell sorting
IRES: internal ribosomal entry site
mdn: median
PQC: protein quality control
PSI: protein stability index
rASA: relative accessible surface area
UPS: ubiquitin-proteasome system
VAMP-seq: variant abundance by massively parallel sequencing.

## Acknowledgements

We acknowledge the use of the FACS, sequencing and computing core facilities at the Department of Biology, University of Copenhagen. We thank Anne-Marie Lauridsen for technical assistance. We thank Prof. Niels Mailand and Dr. Andreas Ingham for assistance with the CRISPR screen. Fig. 1A, Fig. 2A, and Fig. 7 were created with BioRender.com.

## Conflict of interest

K.L.-L. holds stock options in, receives sponsored research from, and is a consultant for Peptone Ltd. The remaining authors have no relevant financial or non-financial interests to disclose.

## Supplementary files

This article includes the following supplementary information.

- Supplementary figures and tables (SupplementaryFigures.pdf).
- Supplementary dataset (SupplementaryFile1.xlsx).

## Data and code availability

All data generated are included in the figures and supplementary files. Sequencing reads are available at the NCBI Sequence Read Archive (SRA) (accession number: PRJNA1359555) and at https://sid.erda.dk/sharelink/BT4LyWUisj. The code is available at GitHub https://github.com/KULL-Centre/_2025_Pedersen_PRKN_BAG6.

## Author contributions

L.P., J.D., I.K.H., E.S.S., M.G.-T., K.E.J., and C.M.H. performed the experiments. L.P., J.D., E.S.S., M.G.-T., V.V., K.E.J., K.L.-L. and R.H.-P. analyzed the data. L.P., K.L.-L, and R.H.-P. conceived the study. L.P. and R.H.-P. wrote the paper.

## Funding

The present work was funded by the Novo Nordisk Foundation challenge programs PRISM (to K.L.-L. & R.H.-P.), REPIN (to R.H.-P.), NNF21OC0071057 (to R.H.-P.), Parkinsonforeningen A2456 (to R.H.-P.) and the Danish Council for Independent Research (Det Frie Forskningsråd) 10.46540/2032-00007B (to R.H.-P.). We acknowledge access to computational resources via a grant from the Carlsberg Foundation (CF21-0392). The funders had no role in study design, data collection and analyses, decision to publish, or preparation of the manuscript.

