## Supplemental figures and tables for "BAG6 and RNF126 are broadly involved in protein quality control of non-native missense protein variants"

### *Supplementary information*


**Supplementary Fig. 1** – *Reduced Parkin abundance in response to geldanamycin.* p.2
**Supplementary Fig. 2** – *Western blotting of knockout cells.* p.3
**Supplementary Fig. 3** – *Complementation in BAG6 and RNF126 knockout cells.* p.4
**Supplementary Fig. 4** – *Flow cytometry of Parkin variants in SGTA knockout cells.* p.5
**Supplementary Fig. 5** – *Parkin variants interact with BAG6.* p.6
**Supplementary Fig. 6** – *Flow cytometry of the Parkin variant library in SGTA knockout cells.* p.7
**Supplementary Fig. 7** – *Overview of low abundance Parkin variants targeted by BAG6.* p.8
**Supplementary Fig. 8** – *The BAG6-responsive variants are buried and destabilized.* p.9
**Supplementary Fig. 9** – *FACS gating strategy on BD FACSymphony for CRISPR screening.* p.10
**Supplementary Fig. 10** – *Flow cytometry gating strategy on the BD FACSJazz instrument.* p.11
**Supplementary Fig. 11** – *Gating strategy for transient co-transfections.* p.12
**Supplementary Fig. 12** – *FACS gating strategy on BD FACSymphony for VAMP seq.* p.13
**Supplementary Table 1** – *ΔPSI scores for pathogenic low abundance Parkin variants.* p.14
**Supplementary Table 2** – *Primers used in this study.* p.15
**Supplementary references** p.16

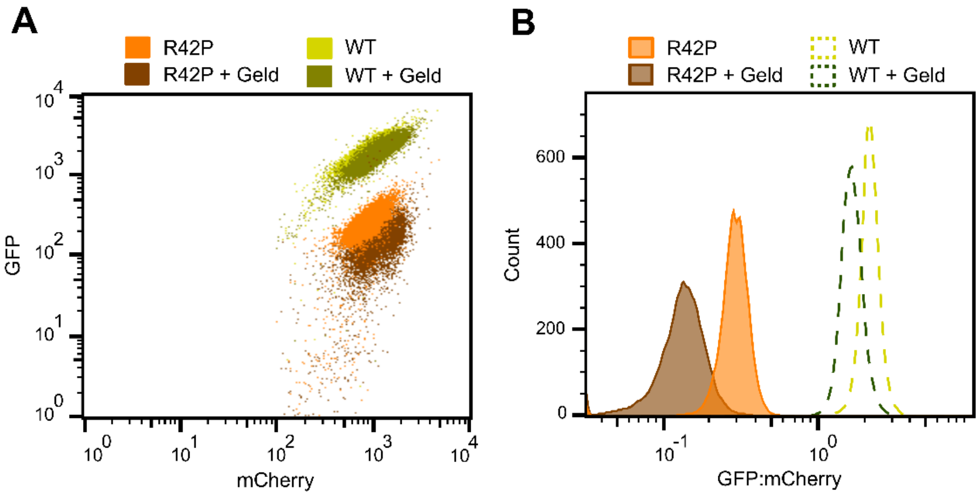


**Supplementary Fig. 1** – *Reduced Parkin abundance in response to geldanamycin.*
Flow cytometry (A) scatter plot and (B) histogram of wild-type Parkin (WT n = $1\times10^4$, WT+Geld n = $9.9\times10^3$) and R42P (R42P n = $1\times10^4$, R42P+Geld n = $1\times10^4$), after the molecular chaperone HSP90 was blocked for 16 hours prior to analyses with 1 μM geldanamycin (+Geld).

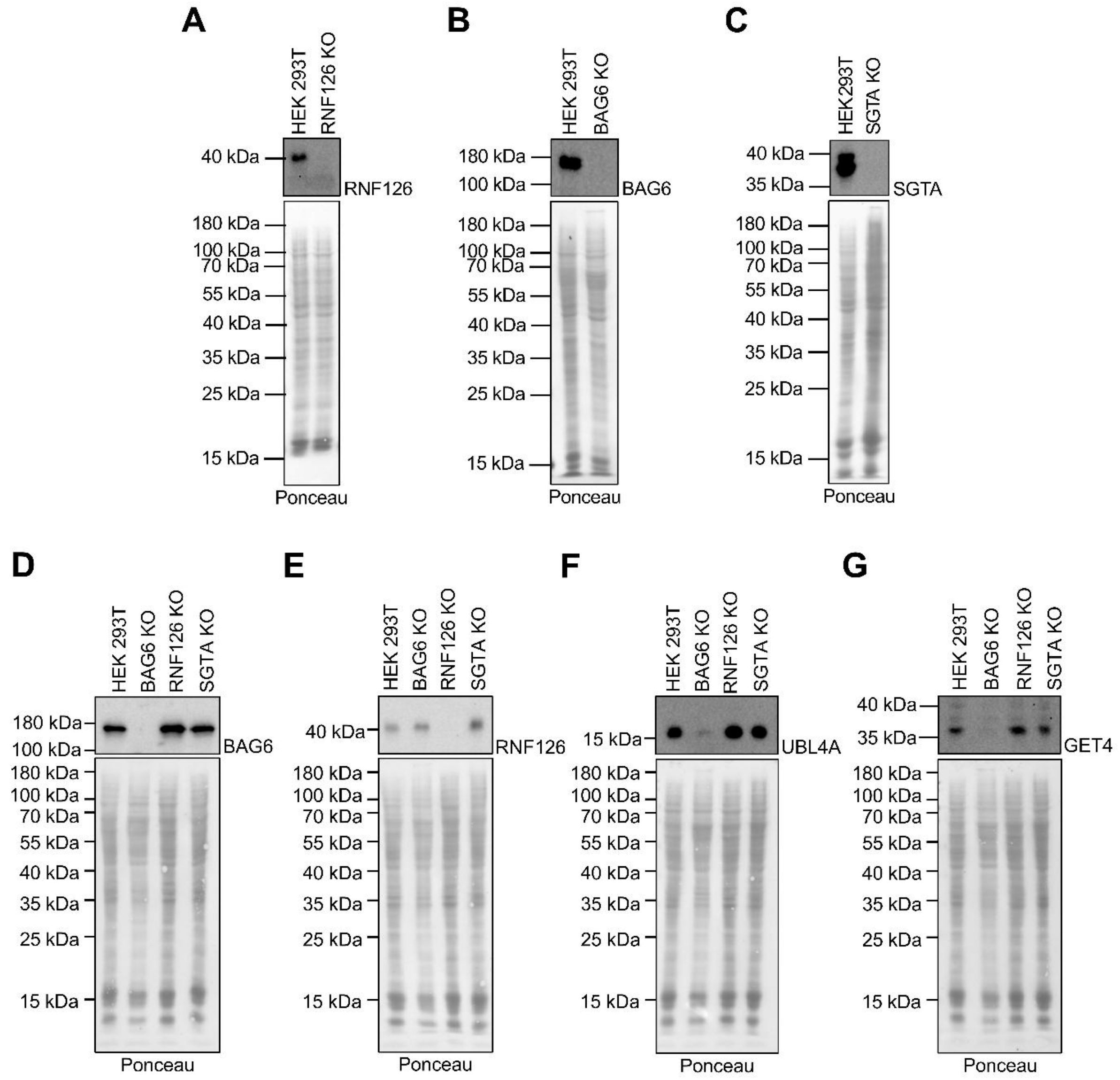


**Supplementary Fig. 2** – *Western blotting of knockout cells.*

Western blotting of whole cell lysates of HEK293T landing pad cells, wild-type (HEK293T) or CRISPR knockouts (KO) for (A) RNF126, (B) BAG6, and (C) SGTA. Ponceau staining was used as a loading control. (D-G) Western blotting of whole cell lysates of HEK293T landing pad cells, wild-type (HEK293T) or the indicated CRISPR knockouts (KO) probed for (D) BAG6, (E) RNF126, (F) UBL4A, and (G) GET4. Ponceau staining was used as a loading control.

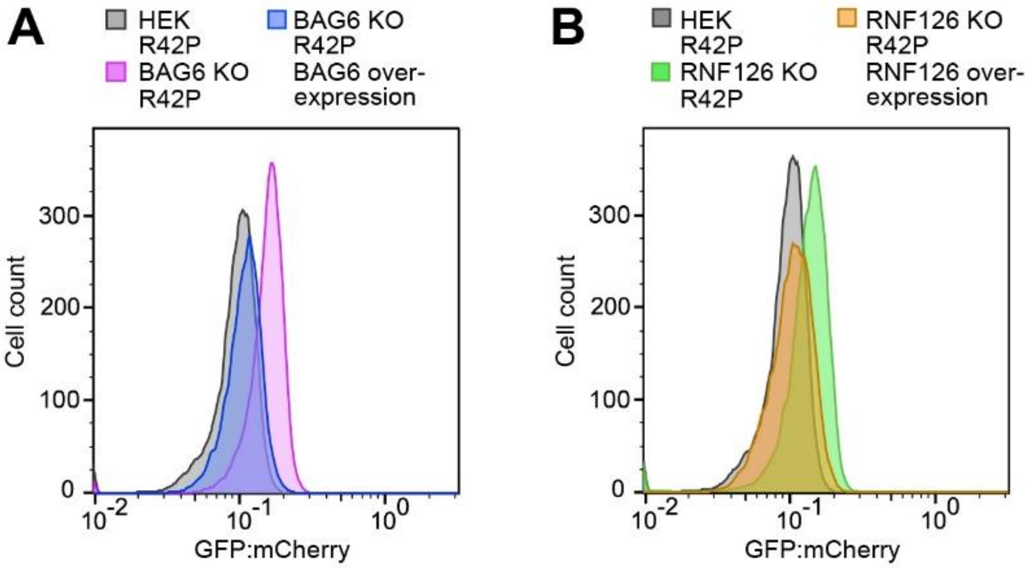


**Supplementary Fig. 3** – *Complementation in BAG6 and RNF126 knockout cells*.
(A) Flow cytometry histograms of Parkin R42P in HEK293T landing pad BAG6 knockout (KO) or (B) RNF126 KO cells. The depleted proteins were re-introduced into the respective KO cell line by transient transfection (HEK R42P (grey, both panels), n=11,662; BAG6 KO R42P (pink), n=11,015; BAG6 KO R42P BAG6 overexpression (blue), n=9,834; RNF126 KO R42P (green), n=11,292; RNF126 KO R42P RNF126 overexpression (orange), n=10,713).

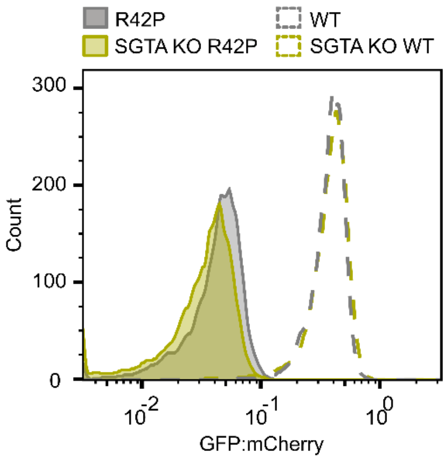


**Supplementary Fig. 4** – *Flow cytometry of Parkin variants in SGTA knockout cells.*
Comparison of flow cytometry histograms of the Parkin wild-type (WT) and R42P in wild-type cells (WT n = $7\times10^3$, R42P n = $7\times10^3$) and SGTA knockout (KO) cells (WT n = $7\times10^3$, R42P n = $7\times10^3$).

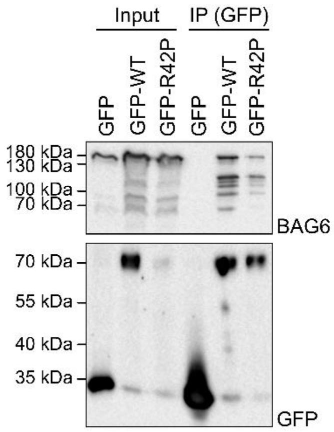


**Supplementary Fig. 5** – *Parkin variants interact with BAG6.*

Wild-type HEK293T landing pad cells expressing either GFP, GFP-tagged wild-type Parkin or GFP-Parkin R42P were lysed (input) and used for immunoprecipitation (IP) using GFP-trap resin. The precipitated material and input samples were separated by SDS-PAGE and analyzed by western blotting using antibodies to BAG6 and to GFP.

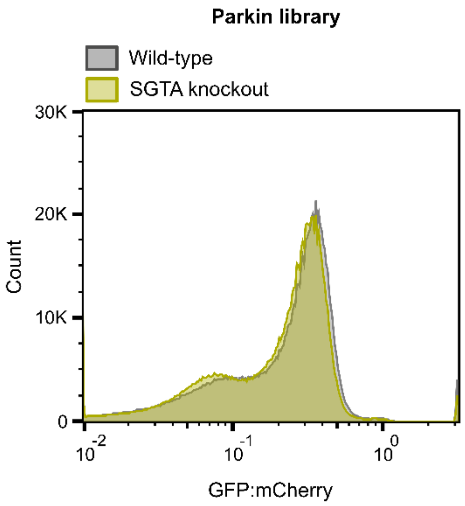


**Supplementary Fig. 6** – *Flow cytometry of the Parkin variant library in SGTA knockout cells.* Comparison of flow cytometry histograms of the Parkin variant library in wild-type cells (grey, n = $8{\times}10^5$) and SGTA knockout cells (yellow, n = $8{\times}10^5$).

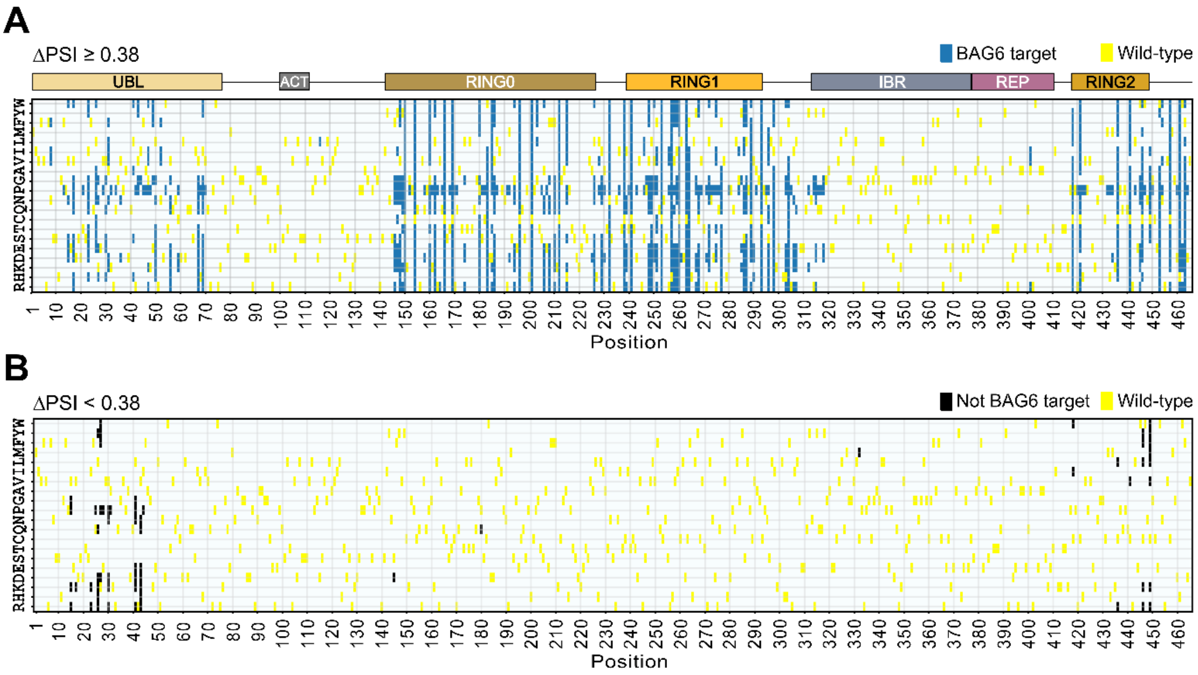


**Supplementary Fig. 7** – *Overview of low abundance Parkin variants targeted by BAG6.*
Heatmaps displaying (A) low abundance Parkin variants that are stabilized in BAG6 knockout cells (blue, ΔPSI ≥ 0.38), and (B) low abundance Parkin variants that are not stabilized in BAG6 knockout cells (black, ΔPSI < 0.38). The wild-type residues are marked yellow. The Parkin domain organization is shown as colored bars.

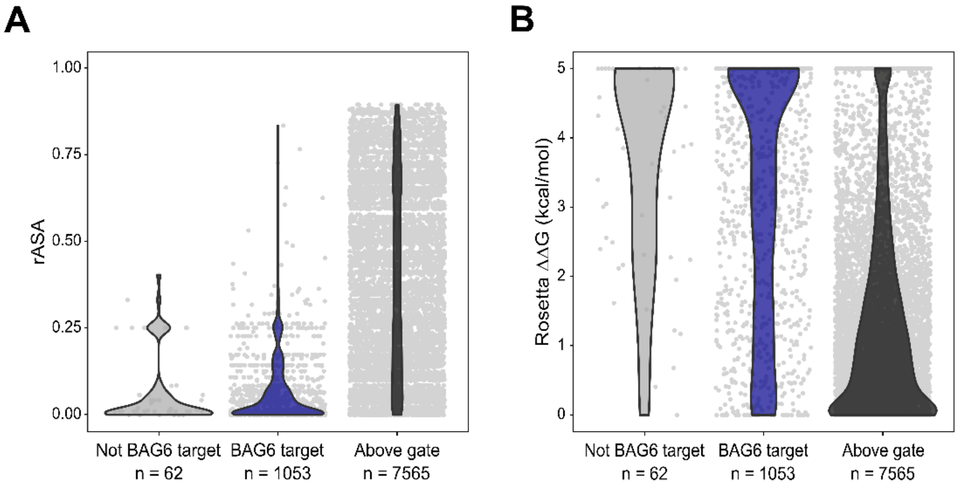


**Supplementary Fig. 8** – *The BAG6-responsive variants are at buried positions and destabilized.* Combined violin and scatter plots showing the (A) relative accessible surface area (rASA) of the variant position and (B) Rosetta predicted destabilization (ΔΔG) of the low abundance Parkin variants that are not (ΔPSI < 0.38) BAG6 targets (grey), low abundance Parkin variants that are (ΔPSI ≥ 0.38) BAG6 targets (blue), and, for comparison, the Parkin variants with higher abundance (above gate, black). The rASA and Rosetta predictions are from Clausen et al. [1].

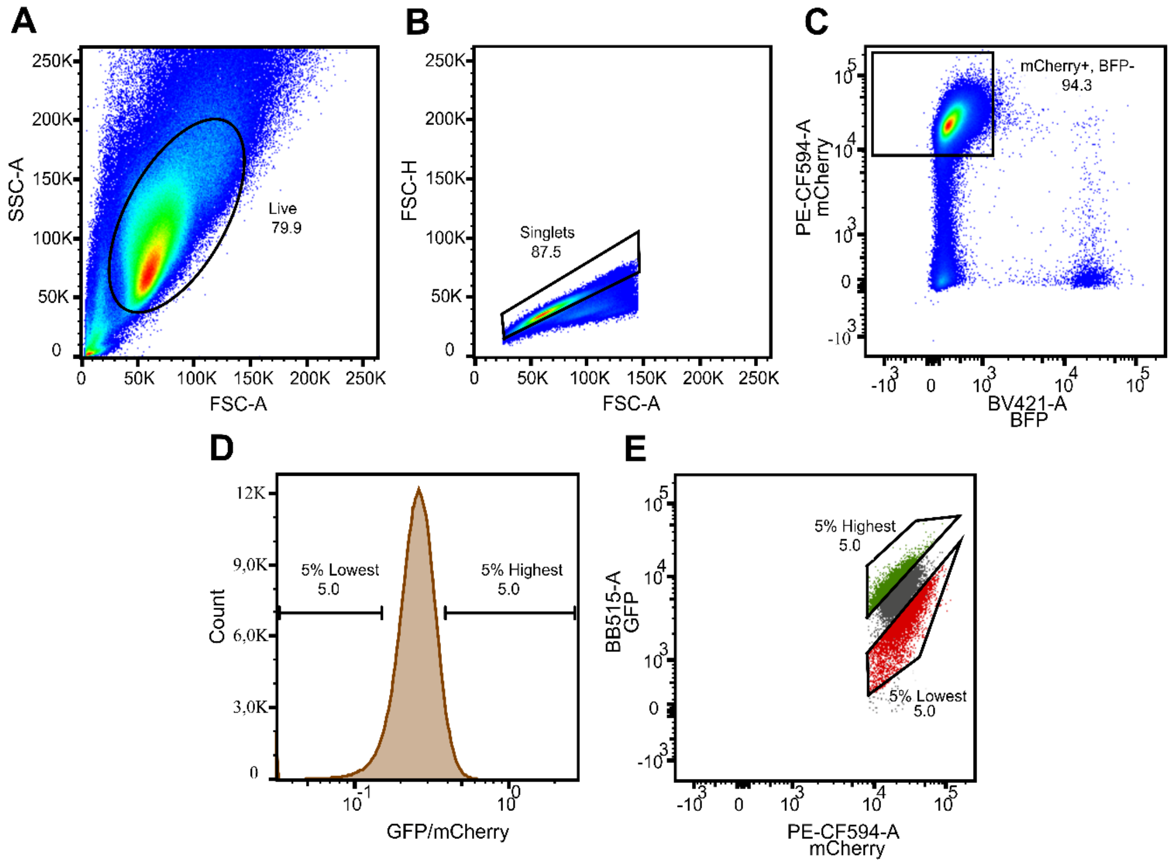


**Supplementary Fig. 9** – *FACS gating strategy on BD FACSymphony for CRISPR screening.*

For fluorescence activated cell sorting (FACS), the cell population was gated as shown for (A) live cells, based on forward and side scattering, (B) singlets, based on the forward scatter height and forward scatter area, and (C) BFP negative and mCherry positive. Based on (D) GFP:mCherry ratios, the cells with the 5% highest and the 5% lowest GFP/mCherry ratios were isolated (E).

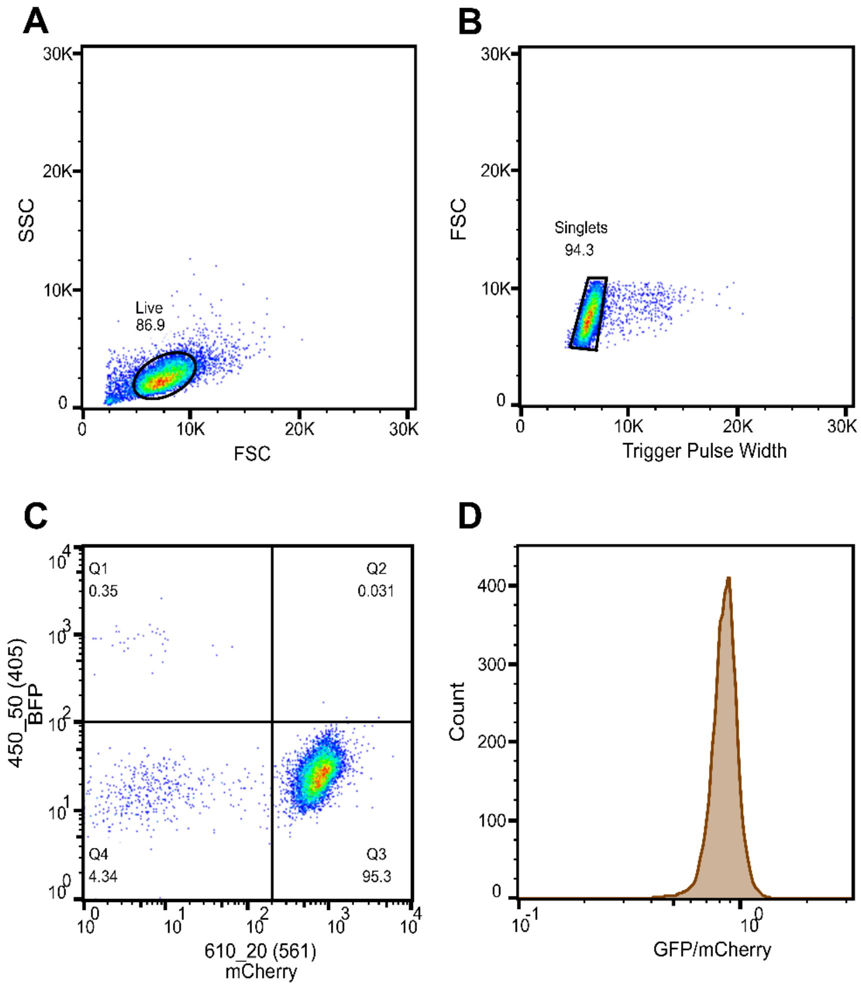


**Supplementary Fig. 10**– *Flow cytometry gating strategy on the BD FACSJazz instrument.*
For determining protein abundance, the cell population was gated as shown for (A) live cells, based on forward and side scattering, (B) singlets, based on the forward scatter and trigger pulse width, and (C) BFP negative and mCherry positive, corresponding to quadrant Q3. Finally, (D) the GFP:mCherry ratios were determined and reported as the shown histogram.

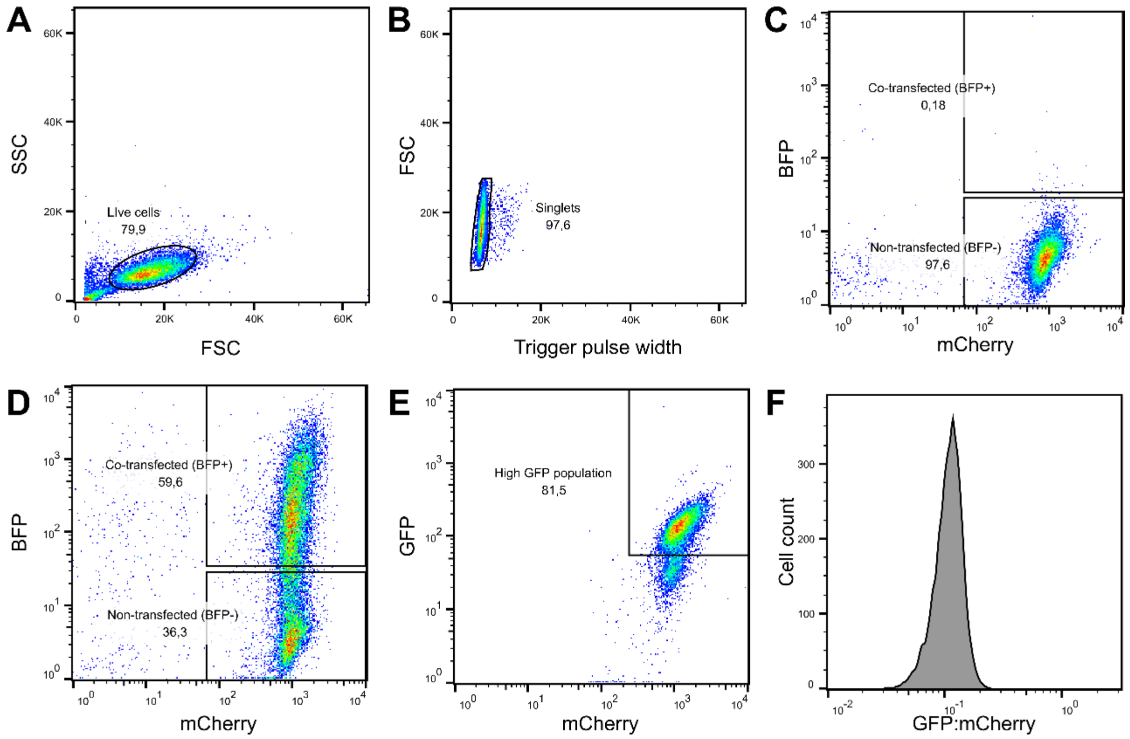


**Supplementary Fig. 11** – *Gating strategy for transient co-transfections.*

(A) Live cells were gated using forward scatter (FSC) and sideward scatter (SSC). (B) Forward scatter (FSC) and trigger pulse width were used to gate for single cells. (C) Cells that had not been co-transfected were BFP negative and mCherry positive, whereas cells that had been transiently co-transfected (D) with BAG6 or RNF126 and BFP were gated as BFP and mCherry positive. (E) Back gating was performed to gate select the main population. (F) Finally, the GFP:mCherry ratios were determined and reported as in the shown histogram.

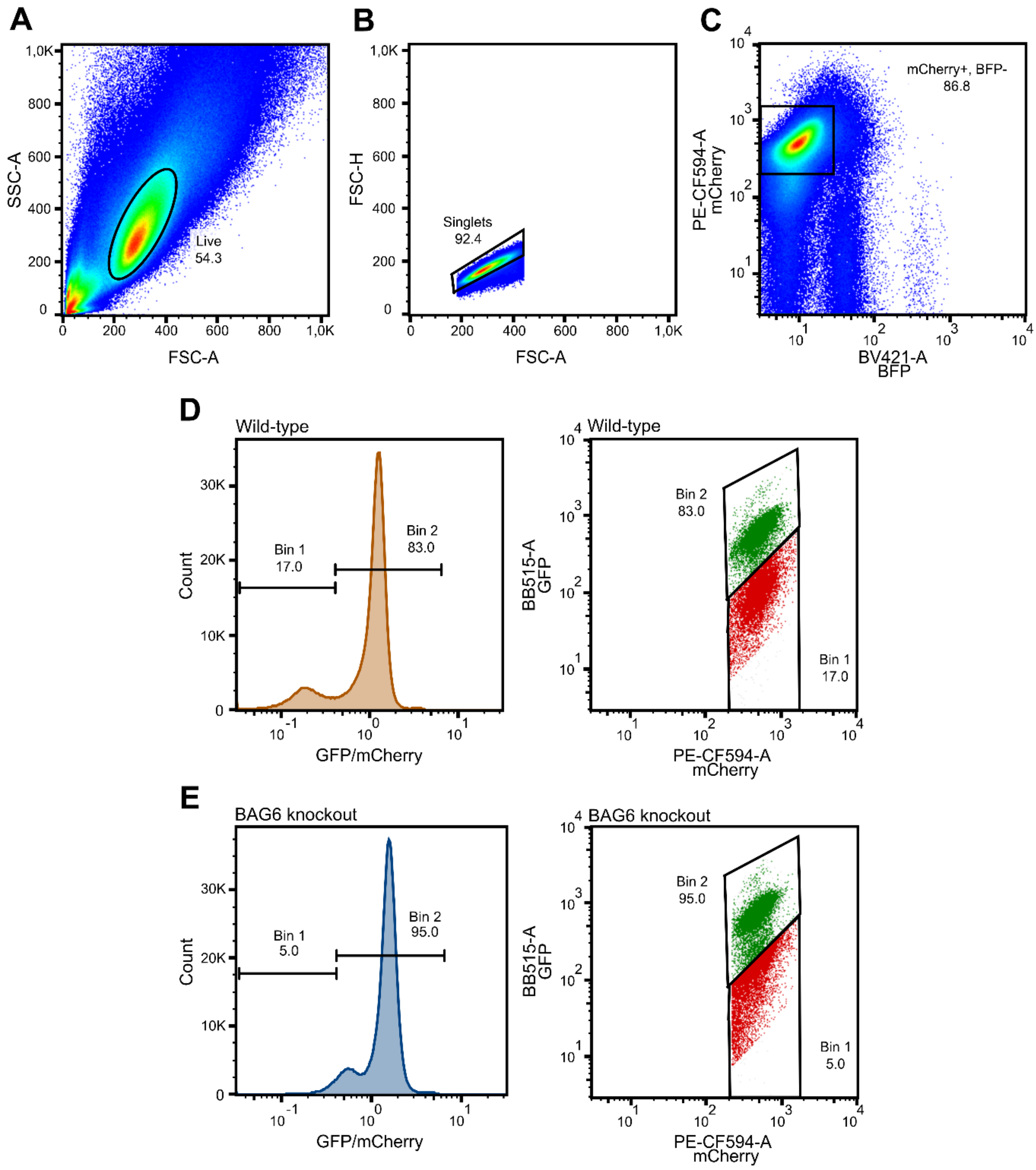


**Supplementary Fig. 12** – *FACS gating strategy on BD FACSymphony for VAMP seq.*
For fluorescence activated cell sorting (FACS), the cell population was gated as shown for (A) live cells, based on forward and side scattering, (B) singlets, based on the forward scatter height and forward scatter area, and (C) BFP negative and mCherry positive. Based on (D) GFP:mCherry ratios, the cells with the 17% lowest GFP:mCherry signal in wild-type cells were isolated. (E) The Parkin variant library in BAG6 knockout cells was sorted using the same gates and bins as the wild-type cells (corresponding to 5% of the library).

**Supplementary Table 1**

*ΔPSI scores for pathogenic low abundance Parkin variants*

| **Pathogenic variant** | **PSI in WT ± SD** | **PSI in BAG KO ± SD** | **ΔPSI** |
|---|---|---|---|
| R42P | 1.044 ± 0.019 | 1.425 ± 0.373 | 0.38 |
| C212Y | 1.009 ± 0.013 | 1.788 ± 0.115 | 0.78 |
| C253Y | 1.045 ± 0.008 | 1.879 ± 0.065 | 0.83 |
| R275W | 1.171 ± 0.054 | 1.959 ± 0.033 | 0.79 |
| C441R | 1.093 ± 0.107 | 1.767 ± 0.023 | 0.67 |

PSI, protein stability index; SD, standard deviation; KO, knockout

**Supplementary Table 2**

*Primers used in this study*

| Primer | Sequence |
|---|---|
| **PARK2index2rev** | ACGCAATTGCAGAACTAGTCCTCATATGTCCTGG |
| **D501_F_PAGE** | AATGATACGGCGACCACCGAGATCTACACTATAGCCTACACTCTTTCCCTACACGACGC TCTTCCGATCTTTGTGGAAAGGACGAAACACCG |
| **D502_F_PAGE** | AATGATACGGCGACCACCGAGATCTACACATAGAGGCACACTCTTTCCCTACACGACGC TCTTCCGATCTATTGTGGAAAGGACGAAACACCG |
| **D701_R_PAGE** | CAAGCAGAAGACGGCATACGAGATCGAGTAATGTGACTGGAGTTCAGACGTGTGCTCTT CCGATCTACTTGCTATTTCTAGCTCTAAAAC |
| **D702_R_PAGE** | CAAGCAGAAGACGGCATACGAGATTCTCCGGAGTGACTGGAGTTCAGACGTGTGCTCTT CCGATCTACTTGCTATTTCTAGCTCTAAAAC |
| **JS_R** | CAAGCAGAAGACGGCATACGAGAT(NNNNNNNN)GGGTTAGCAAGTGGCAGCCT |
| **LC1020** | CCAGGACATATGAGGACTAG |
| **LC1031** | GGGTTAGCAAGTGGCAGCCTTCTCCTTAATCAGCTCTTCG |
| **LC1040** | AAGAACCGCTAGAAGCGTCGCTGTACAAATAGTT |
| **LC1041** | CGAGAAAGCTAGCGCAAACGACTACTCGCA |
| **LC1042** | CTGATTAAGGAGAAGGCTGCCACTTGCTAACCC |
| **LCV2 fwd** | GAGGGCCTATTTCCCATGATTC |
| **LCV2 rev** | GTTGCGAAAAAGAACGTTCACGG |
| **PCR2_FW** | AATGATACGGCGACCACCGAGATCTACAC(NNNNNNNN)CCAGGACATATGAGGACTAG |
